# Dramatically reduced spliceosome, intronome, and splicing efficiency in *Cyanidiococcus yangmingshanensis* and *Cyanidium caldarium*

**DOI:** 10.64898/2026.05.17.725761

**Authors:** Viktor A. Šlat, Martha R. Stark, Stephen D. Rader

## Abstract

Eukaryotic pre-mRNA splicing is catalyzed by the spliceosome, whose ribonucleoprotein composition and the number of intron substrates it acts upon vary widely across eukaryotic lineages. The red alga *Cyanidioschyzon merolae* possesses a reduced spliceosome lacking the U1 snRNP, and an unusually small intron repertoire. We asked whether these traits are unique to *C. merolae* or shared across the related Cyanidiales and Cyanidioschyzonales lineages, as well as how they relate to splicing efficiency under light conditions relevant to photosynthetic growth. Genomic and transcriptomic analysis of *C. merolae*, *Cyanidiococcus yangmingshanensis*, and *Cyanidium caldarium* reveal that all three species harbour a reduced, but broadly conserved, set of splicing proteins. Strikingly, covariance model searches failed to detect U1 snRNA in either *C. yangmingshanensis* or *C. caldarium*, establishing U1 loss as a shared feature of all three lineages. We identified only 39 introns in *C. merolae*, 40 in *C. yangmingshanensis*, and 54 in *C. caldarium*. Splicing efficiencies were 42–50%, substantially lower than most organisms in which splicing has been measured, but low splicing is compensated by 2–4× higher expression of intron-containing genes than intron-lacking genes. Notably, light can enhance splicing efficiency in *C. merolae* and *C. yangmingshanensis* by up to 100%. Furthermore, the splice site and branch site consensus sequences are highly conserved and similar to those found in hemiascomycetous yeasts such as *Saccharomyces cerevisiae*. 85% of introns contain an in-frame stop codon with a strong bias towards the 5′ end of the intron. These results indicate that dramatic streamlining of the spliceosome and intronome, together with inefficient splicing, predated the divergence of these lineages ∼320 million years ago, and is therefore a defining molecular trait of these extremophilic red algae.

## Introduction

Discovered in the latter half of the 1970s (Berget et al. 1977; Chow et al. 1977), the eukaryotic process of precursor messenger RNA (pre-mRNA) splicing involves the removal of intron sequences and the ligation of the remaining exon sequences (Will and Luhrmann 2011; Matera and Wang 2014; Wan et al. 2020). This reaction is catalyzed by the spliceosome, a dynamic, multi-megadalton ribonucleoprotein complex composed of five small nuclear RNAs (snRNAs), U1, U2, U4, U5, and U6, as well as a large number of associated proteins. Traditional model organisms for the study of splicing include *Homo sapiens*, with more than 300 splicing-associated proteins (Rappsilber et al. 2002) acting on over 200,000 introns (Sakharkar et al. 2004), and the budding yeast *Saccharomyces cerevisiae*, with approximately 100 splicing proteins (Fabrizio et al. 2009) acting on fewer than 300 introns (Parenteau et al. 2008). However, the splicing landscapes of many eukaryotic lineages remain poorly characterized, and extending splicing studies beyond traditionally studied organisms is essential for understanding how the spliceosome evolved, which cannot be inferred from a few model systems alone.

One such lineage is the unicellular red algal class Cyanidiophyceae, the most ancient of the seven classes comprising the red algal phylum Rhodophyta, having diverged approximately 1.5 billion years ago (Yoon et al. 2004). Cyanidiophyceae is composed of the thermoacidophilic orders Galdieriales, Cyanidiales, and Cyanidioschyzonales, whose members can grow at temperatures as high as 65 °C and at pH values as low as 0, as well as the mesophilic order Cavernulicolales, whose members prefer temperatures of 18–25 °C and pH values of 5–8, reflecting a broad ecological and physiological diversity within an ancient evolutionary framework (Park et al. 2023; Yoon and Del Guacchio 2024). Molecular clock estimates place the earliest divergence within Cyanidiophyceae at approximately 780 million years ago for the Galdieriales, followed by Cavernulicolales at 560 million years ago, and most recently the split between Cyanidiales and Cyanidioschyzonales at 320 million years ago (Yang et al. 2016). Focusing on the thermoacidophilic orders, the Galdieriales consists of a single genus, *Galdieria*, which includes species such as *G. sulphuraria* and *G. yellowstonensis*; the Cyanidiales comprises a single genus and species, *Cyanidium caldarium*; and the Cyanidioschyzonales includes two genera, *Cyanidioschyzon* and *Cyanidiococcus*, represented by the species *C. merolae* and *C. yangmingshanensis*, respectively (Park et al. 2023). Hereafter, we refer to Cyanidiales and Cyanidioschyzonales collectively as Cx, and to the species *C. caldarium*, *C. merolae*, and *C. yangmingshanensis* as Cc, Cm, and Cy, respectively.

Cm represents an extreme outcome of spliceosome and intron evolution. An established model organism for cell biology (Miyagishima and Tanaka 2021), and an emerging one for pre-mRNA splicing (Stark et al. 2015; Reimer et al. 2017; Schärfen et al. 2022; Black et al. 2023; Wong et al. 2023), Cm has a dramatically reduced complement of about 50 core splicing proteins inefficiently splicing fewer than 40 introns, as well as the striking absence of the U1 snRNA and its associated proteins (Stark et al. 2015; Reimer et al. 2017; Schärfen et al. 2022; Wong et al. 2023). In sharp contrast, the last common ancestor of the Rhodophyta is estimated to have possessed more than 1,600 introns and more than 150 splicing proteins, highlighting the massive loss of introns and splicing proteins in Cm over the course of evolution (Bhattacharya et al. 2018). By comparison, the related thermoacidophile *G. sulphuraria* retains the full complement of snRNAs and more than 140 splicing proteins, efficiently splicing over 13,000 introns (Qiu et al. 2018; Wong et al. 2023).

These extremes raise compelling questions about the evolutionary plasticity and minimal requirements of the spliceosome. In particular, whether the absence of the U1 small nuclear ribonucleoprotein (snRNP), a key component of the spliceosome, and the parallel loss of splicing proteins and introns, are unique to Cm. By examining red algal strains that diverged hundreds of millions of years ago, we sought to determine whether spliceosome reduction and intron loss reflect a species-specific anomaly or a shared evolutionary feature, and to investigate the evolutionary limits of spliceosomal streamlining. To this end, we leveraged newly available genomic and transcriptomic data from one strain of Cc and two equivalent strains of Cy (Liu et al. 2020; Cho et al. 2023) to conduct a comprehensive comparative bioinformatic analysis.

In this study, we demonstrate that the U1 snRNA and its associated proteins are absent not only from Cm but also from Cc and Cy, while a broadly conserved set of approximately 70 splicing proteins is retained across all three species. This establishes U1 loss as a shared, lineage-defining feature of the Cx rather than a peculiarity of Cm, and demonstrates that spliceosome reduction predates the divergence of these three lineages. Furthermore, the small number of introns that persist in these species are spliced with relatively low efficiency, indicating that spliceosome reduction and intron loss are accompanied by altered splicing performance.

## Results

### Genomic features, chromosomal synteny, and average amino acid identity

To ensure an accurate foundational interpretive context, we verified the core nuclear genomic features of the red algal species examined in this study. The genome of Cm strain 10D is the largest among the Cx species analyzed, measuring 16.6 Mbp with a GC content of 55%. Two Cy genomes have been published, strain 8.1.23 F7 and THAL066, with sizes of 12.0 Mbp and 12.1 Mbp, respectively, and a GC content of 55% (Liu et al. 2020; Cho et al. 2023). In contrast, Cc strain 063 E5 has the smallest genome at 8.8 Mbp and exhibits the highest GC content at 66%. The nuclear genomes of the Cx members Cm, Cy, and Cc are each organized into 20 chromosomes. In comparison, *G. yellowstonensis* strain 108.79 E11, a member of the more distant Galdieriales and the most complete genome assembly currently available for that lineage, has a genome size of 14.5 Mbp spread across 76 scaffolds and a much lower GC content of 40%.

To investigate the extent of genome conservation and divergence among the members of the Cx, we compared chromosomal synteny across Cm, Cy, and Cc, as visualized in Fig. 1, which shows conserved syntenic blocks and reveals patterns of genome rearrangement that inform their overall syntenic relationships. Examination of these syntenic relationships demonstrates a high level of structural conservation between Cm and Cy (Supplemental Fig. S1), reflecting a recent common ancestry that has largely preserved genomic integrity. In contrast, the genome of Cc displays greater chromosomal rearrangement and genomic divergence compared to both Cm (Supplemental Fig. S2) and Cy (Supplemental Fig. S3), reflecting a more distant common ancestry. No substantial differences in chromosomal synteny were observed between the 8.1.23 F7 and THAL066 strains of Cy (Supplemental Fig. S4). Moreover, the average pairwise identity of orthologous intronic sequences in the two strains of Cy was 99%, ranging from 97% to 100%, indicative of the near identity and equivalence of the strains. The patterns of chromosomal synteny observed across the Cx (Fig. 1) are in line with those previously reported (Liu et al. 2020; Cho et al. 2023).

**Figure 1.**
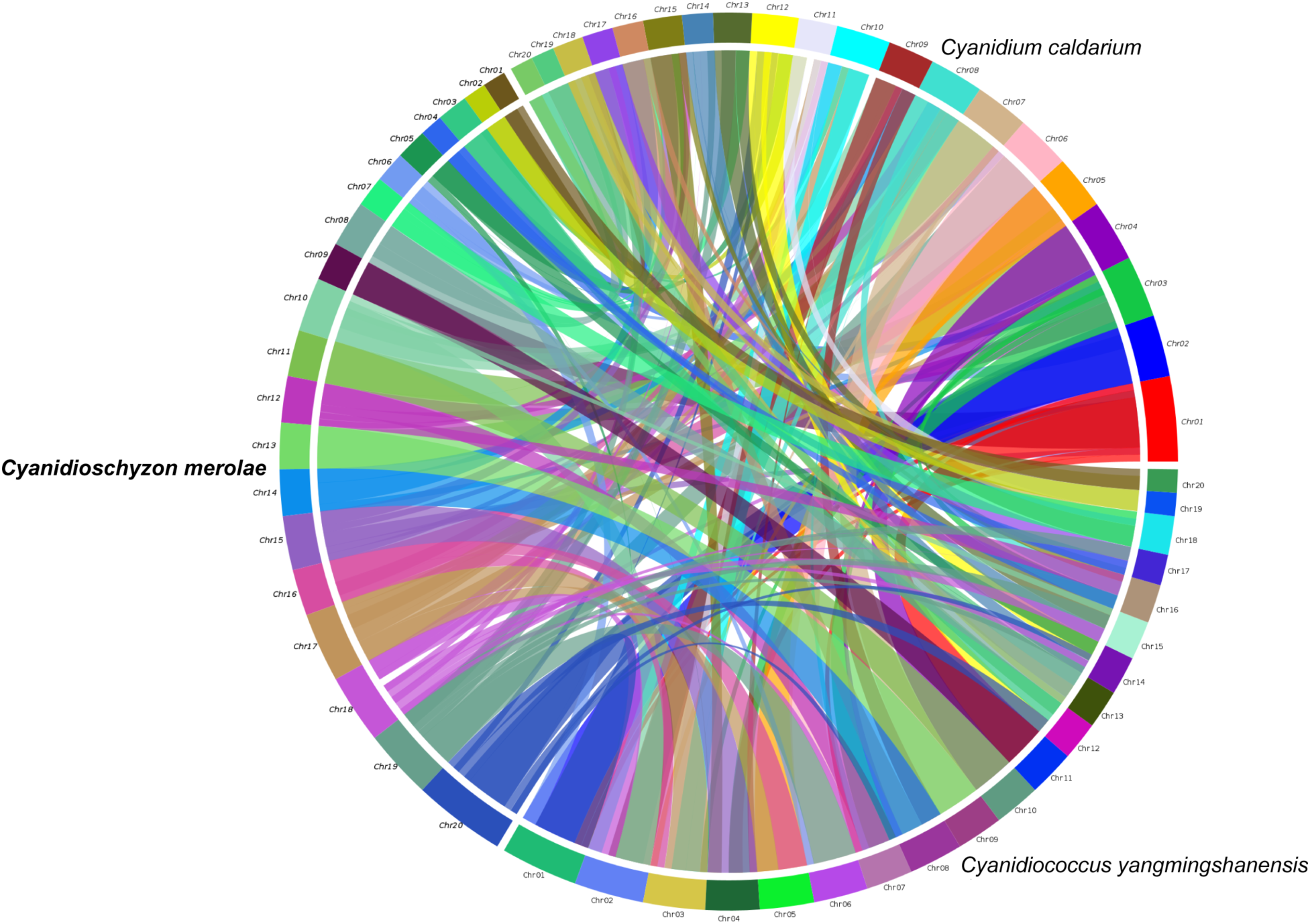
Chromosomal synteny across the Cx. All nuclear chromosomes are shown for *C. merolae* 10D, *C. yangmingshanensis* 8.1.23 F7, and *C. caldarium* 063 E5. Each species has been allocated a third of the circle. Arcs along the circumference of the circle represent chromosomes, while connecting arcs within the circle represent synteny blocks whose colour is determined by the chromosome of the species whose name appears in bold.

To further assess the evolutionary relationships among the Cx, as well as the fellow red alga *G. yellowstonensis*, we performed an average amino acid identity (AAI) analysis (Fig. 2) so as to quantify genomic relatedness at the protein level. The highest AAI value of 65% was observed between Cm and Cy, indicating the closest evolutionary relationship among the species analyzed. Intermediate AAI values were obtained when comparing Cc to Cy and Cm, with values of 52% and 52%, respectively, suggesting a moderately distant evolutionary relationship. In contrast, the lowest AAI values were found when comparing *G. yellowstonensis* to Cc, Cm, and Cy, with values of 44%, for each, consistent with its position as the most distantly related species in the group. The computed AAI values (Fig. 2) accord with the divergence times reported previously (Yang et al. 2016).

**Figure 2.**
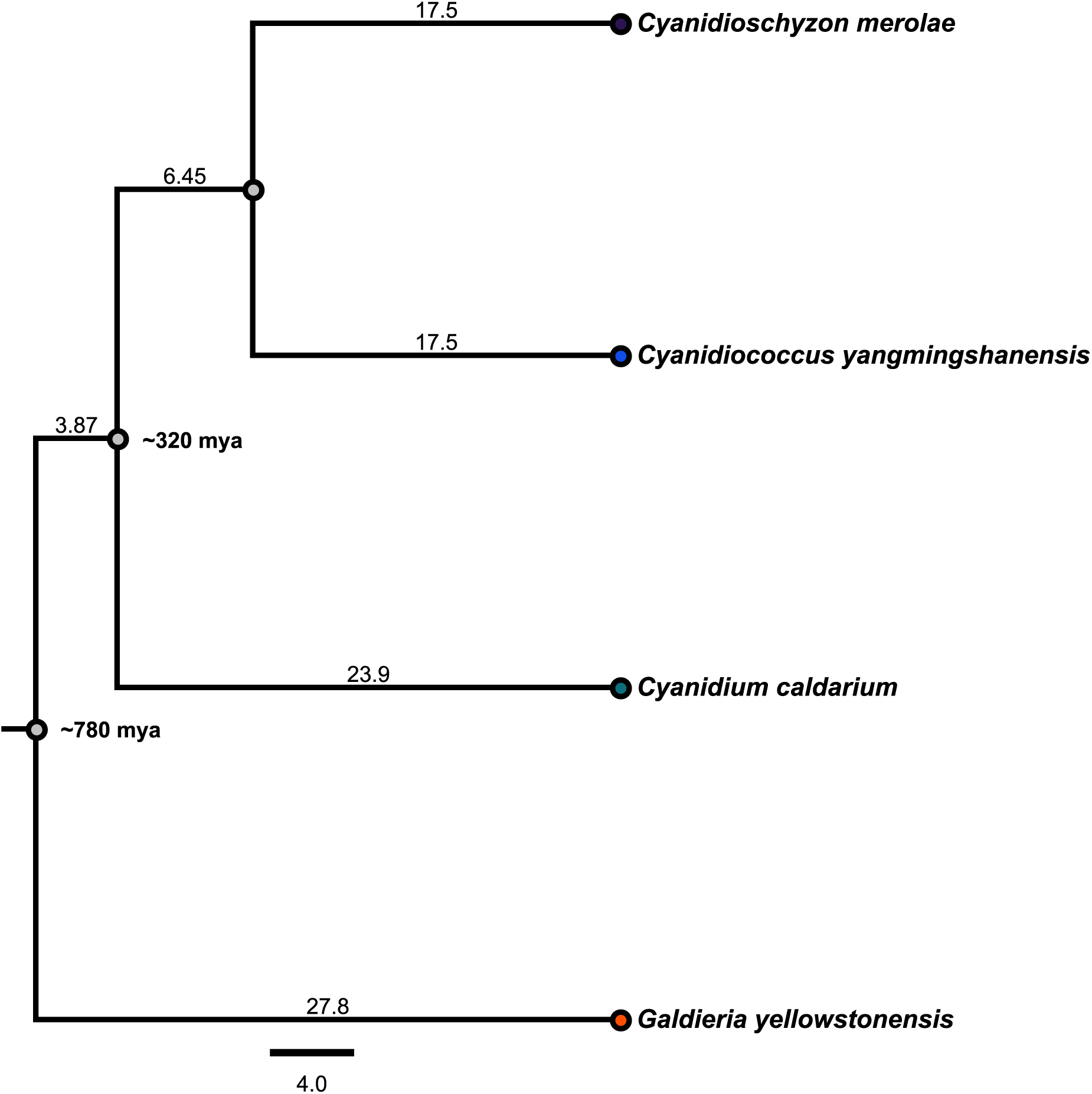
Phylogenetic tree of the Cx. Branch lengths and scale show average amino acid identity (%) values. The outgroup is a member of the Galdieriales. Divergence times are shown in million years ago (mya) and are based on molecular clock estimates (Yang et al. 2016).

### Identification of splicing proteins

We previously reported that Cm has a substantially reduced complement of splicing proteins (Stark et al. 2015; Reimer et al. 2017; Black et al. 2023) and sought to ascertain whether Cy and Cc possess a similar degree of reduction. We identified the deemed orthologs of every Cm splicing protein in Cy and Cc (Supplemental Table S1) by employing a reciprocal best hits strategy as a proxy for orthology (Ward and Moreno-Hagelsieb 2014). Consistent with the previously established absence of U1-associated proteins in Cm, none were identified in either Cy or Cc. Interestingly, five core splicing proteins are encoded by multiple paralogous genes in Cm, Cy, or both, whereas each is represented by a single gene in Cc, indicating limited lineage-specific gene duplication within an otherwise streamlined spliceosome.

To better understand the conservation or divergence of the remaining splicing proteins, we analyzed pairwise identity over full-length sequences (Fig. 3). A small subset of the 65 splicing proteins consistently showed markedly elevated conservation across all three species: Cbc2 (NCBP2), Dhh1 (DDX6), Fal1 (EIF4A3), LSm1 (LSM1), LSm2 (LSM2), LSm3 (LSM3), LSm6 (LSM6), Prp43 (DHX15), Snu13 (SNU13), and Sub2 (DDX39B). These are all notable for having established functions outside of splicing. Otherwise, most splicing proteins exhibited below-average conservation relative to genome-wide AAI. Analysis of the orthologous splicing proteins revealed that the Cx share conserved protein length and electrostatic properties, but Cc is distinguished by a shifted amino acid composition, including higher GC-rich and hydrophobic residue content (Supplemental Note 1; Supplemental Fig. S5). These measurements expose substantial heterogeneity in evolutionary conservation (Fig. 3).

**Figure 3.**
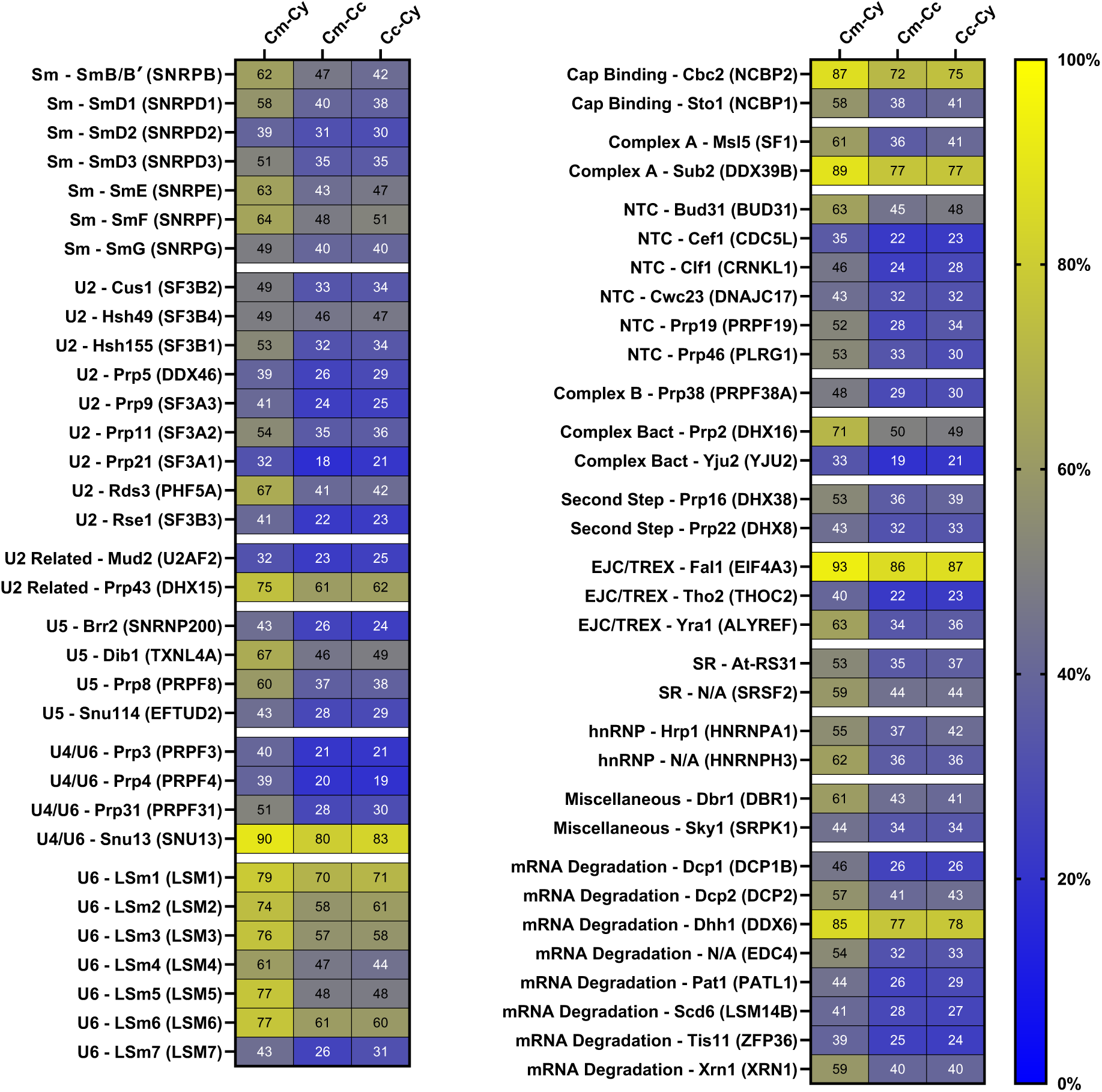
Heatmap of orthologous splicing proteins in the Cx. Values indicate percent pairwise identity across the complete length of each protein. Each protein is categorized by particle or step followed by its name in *S. cerevisiae* and *H. sapiens* (At refers to *A. thaliana*). Paralogs have been omitted from the heatmap.

Building on our earlier observation that several Cm splicing proteins lack conserved domains and instead contain novel sequence features (Black et al. 2023), we extended this analysis to the broader Cx. We examined sequence alignments and domain architectures (Supplemental Fig. S6), identifying the absence of domains and regions conserved in the well-annotated model eukaryotes *H. sapiens* or *S. cerevisiae*. In total, 19 Cx splicing proteins lack one or more domains or regions present in their human or yeast counterparts (Supplemental Table S2), and 6 harbour novel sequence regions (Supplemental Table S3), indicating extensive and largely conserved remodelling of splicing proteins that likely predates divergence within the Cx (Supplemental Note 2).

To assess whether any splicing proteins are present in Cy or Cc that may be absent in Cm, we applied a comprehensive reciprocal best hits strategy using all annotated splicing protein sequences from *Arabidopsis thaliana*, *H. sapiens*, *S. cerevisiae*, and *Schizosaccharomyces pombe*. We identified two additional splicing protein candidates in Cy (SR proteins LDC2 and RSZ22A) and four in Cc (Cwc24, U11/U12 snRNP 31 kDa, RS2Z32, and PT-WD) relative to Cm (Supplemental Table S4; Supplemental Fig. S7).

### Identification of snRNAs

We previously reported the unexpected absence of U1 snRNA in Cm (Stark et al. 2015). To ascertain whether this peculiarity is unique to Cm or shared amongst the broader Cx, we assessed the presence and conservation of the spliceosomal snRNAs in Cy and Cc. Covariance model searches identified the U2, U4, U5, and U6 snRNAs in both species, but failed to detect a U1 snRNA in either, establishing U1 loss as a conserved feature of the Cx (Supplemental Table S5).

To assess conservation of the retained snRNAs, we aligned their sequences, revealing strong conservation of core functional elements across the Cx, including the U2 branchpoint-binding region, the U5 invariant loop 1, the U6 ACAGA-box motif, and the Sm- and LSm-binding sites (Supplemental Fig. S8). U2, U5, and U6 snRNAs are similar in length, whereas the Cc U4 snRNA is 75% longer due to an extended 3′ end. The U2 and U6 snRNAs are the most conserved (69% average identity), followed by the U4 and U5 snRNAs (46% average identity). The Cc snRNAs possess a higher average GC content of 61% compared to Cm and Cy (51%), with a greater predicted thermodynamic stability than those of Cm and Cy (an average free energy value of -562 kJ/mol compared to -395 kJ/mol).

Predicted secondary structures further support this pattern of conserved function with lineage-specific remodelling. U2 snRNA structures are highly conserved across Cm (Supplemental Fig. S9), Cy (Fig. 4), and Cc (Fig. 5). While the U4, U5, and U6 snRNAs for Cm and Cy are highly conserved, those of Cc exhibit pronounced structural divergence: Cc U4 possesses a shorter stem III, a longer stem IV, and a uniquely prominent, 158-nt hairpin structure in place of stem V; Cc U5 core and 3′ regions are fairly conserved, but the structure of the 5′ region has markedly diverged, replacing the long-range stem found in Cm and Cy with a short-range hairpin (Figs. 4 and 5); Cc U6 has a shorter 5′ stem-loop, immediately downstream of which is a small hairpin structure that is unique to Cc. Together, these data indicate strong conservation of essential snRNA features alongside selective structural diversification within the Cx.

**Figure 4.**
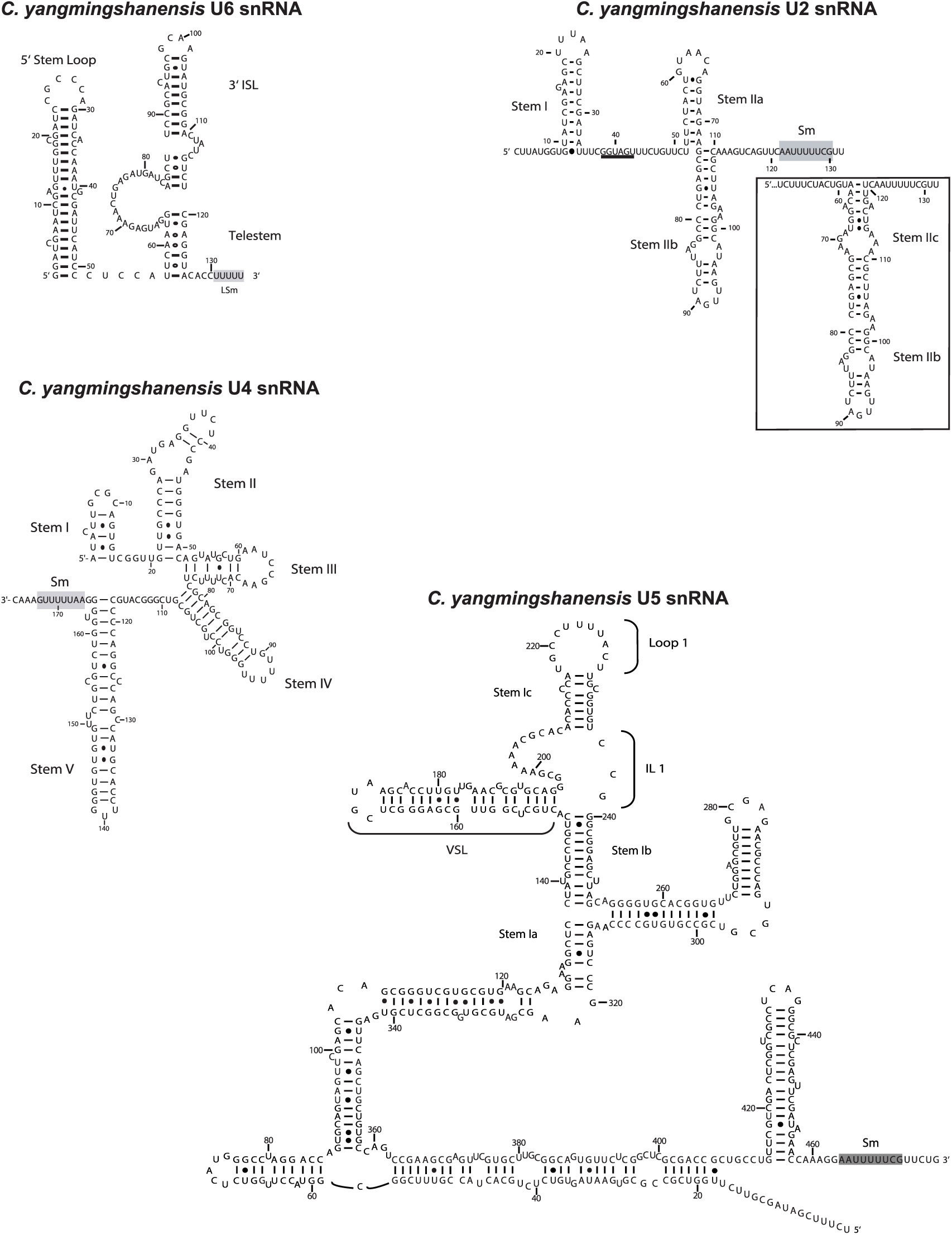
Predicted secondary structures of *C. yangmingshanensis* 8.1.23 F7 snRNAs. The inset shows the alternative U2 toggle structure, in which stem IIc replaces stem IIa. The U2 branchpoint-binding region is underlined, and the Sm- and LSm-binding sites are highlighted in grey.

**Figure 5.**
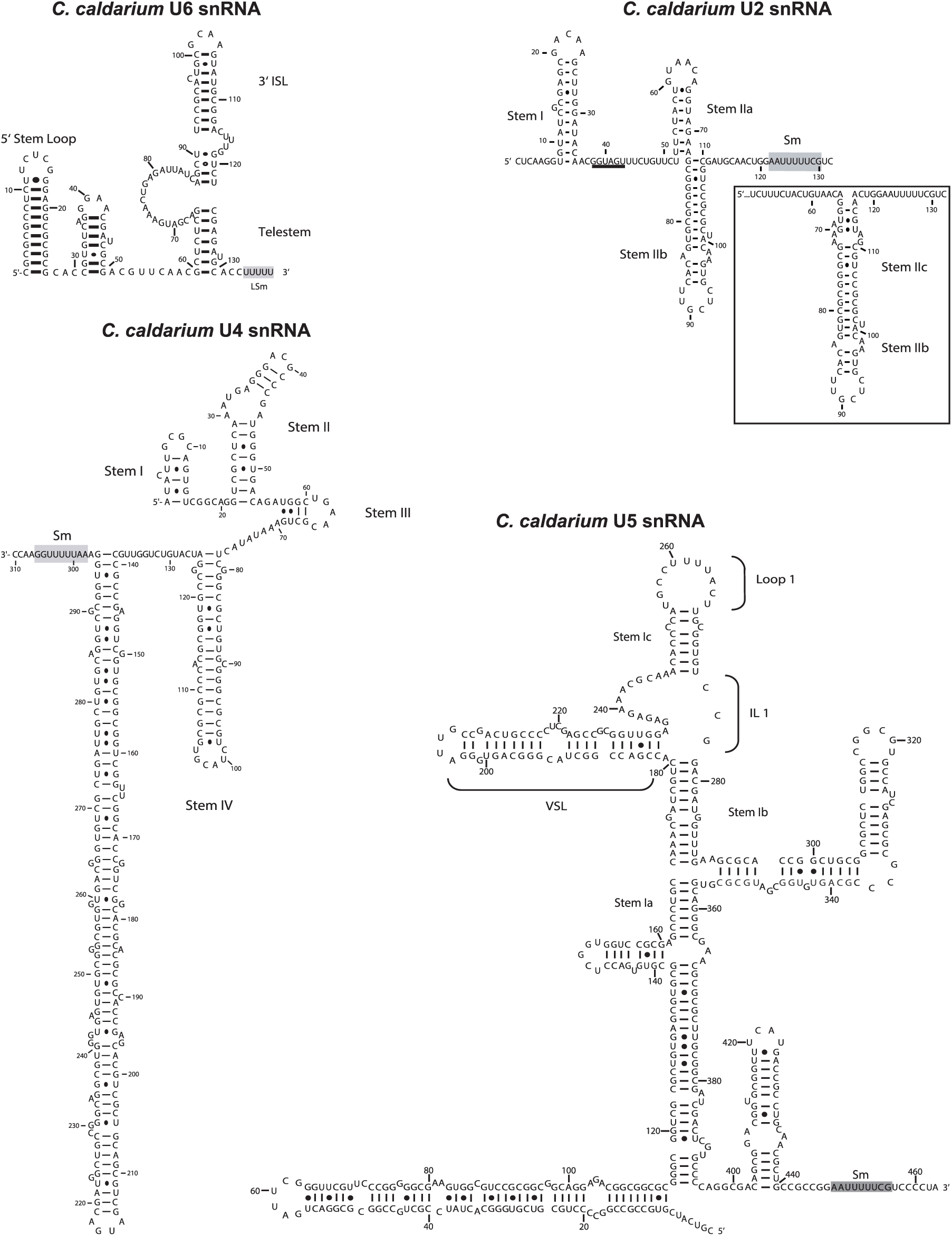
Predicted secondary structures of *C. caldarium* 063 E5 snRNAs. The inset shows the alternative U2 toggle structure, in which stem IIc replaces stem IIa. The U2 branchpoint-binding region is underlined, and the Sm- and LSm-binding sites are highlighted in grey.

### Identification of intron-containing genes

We previously reported that Cm harbours an exceptionally small number of intron-containing genes (ICGs) (Stark et al. 2015; Wong et al. 2023). Extending this analysis to Cy and Cc, we found that all three species possess similarly reduced intronomes. We identified 38 ICGs in Cm, hosting a total of 39 introns, of which 26 were previously annotated, 1 was misannotated, and 12 are novel compared to the current NCBI annotation (Supplemental Table S6; Supplemental Fig. S10A). An intron is considered to be misannotated if the coordinates for one or both of its splice sites were found to be incorrect. In our previous work (Wong et al. 2023) employing a different methodology, we identified all but one of these novel introns, with CmICG25 (CMQ163T) having escaped detection due to its unique 5′ splice site dinucleotide of ‘GA’. Similarly, there were 38 ICGs identified in Cy, hosting a total of 40 introns, of which 32 are annotated, 3 are misannotated, and 5 are novel (Supplemental Table S7; Supplemental Fig. S10A). Fifty ICGs were identified in Cc, hosting a total of 54 introns, of which 38 are annotated, 2 are misannotated, and 14 are novel (Supplemental Table S8; Supplemental Fig. S10A). A complete list of the orthologous ICGs found in the Cx, together with associated coding sequence corrections and discoveries, is provided in Supplemental Table S9. Cx introns are predominantly found interrupting coding sequences, though some can be within the 5′ untranslated region (5′ UTR), 3′ untranslated region (3′ UTR), or even a noncoding RNA (ncRNA) (Supplemental Fig. S10B).

We previously observed that splicing in Cm is inefficient (Schärfen et al. 2022), raising the question of whether unspliced transcripts could be translated into functional proteins. To address this, we looked at the distribution of termination codons in phase with exon1 (or exon2 for the rare two-intron transcripts). We found that the vast majority (85%) of Cx introns interrupting coding sequences contained a premature termination codon (PTC), whose relative position within each intron revealed a pronounced 5′ bias (Supplemental Fig. S10F). Consistent with this 5′ bias, 68% of PTC-containing introns had a PTC within the first quarter of the intron. To rule out potential systematic intron prediction error, we tested whether intron lengths were evenly distributed modulo three, and found no significant deviation from expectation. In contrast, intron phase distributions showed phase 2 introns to be most prevalent across the Cx compared with the phase 0 predominance typical of other eukaryotes (Supplemental Note 3; Supplemental Fig. S10C,D). We analyzed the relative positions of introns within protein-coding sequences, normalized from the 5′ end (position 0) to the 3′ end (position 1) of each coding sequence, and observed that the Cx exhibit a mild 5′ intron bias (Supplemental Fig. S10E).

Given the limited divergence of the Cx splicing systems, we asked whether sequence conservation in ICGs takes place at the protein level (amino acid identity), or at the DNA level (coding sequence identity), using intron identity as a comparator. While the range of identity was fairly large, amino acid identity was most conserved across the Cx ICGs, followed by coding sequence identity, and intron identity (Fig. 6). Although this overall trend in identity was consistent with expectations, it is noteworthy that at the level of individual ICGs intriguing exceptions emerge. There are multiple instances in which specific proteins are remarkably well-conserved (e.g. CmICG30/CMS262C, a ribosomal protein gene). Interestingly, for certain orthologous ICGs, the level of conservation in intron sequences not only approaches, but in some instances exceeds, that of the associated coding sequences (e.g. CmICG08/CMG006C or CmICG21/CM0159C). Across the ICGs, protein lengths were conserved among the Cx, but Cc showed significantly higher coding sequence GC content and a corresponding enrichment of GC-rich amino acids (Supplemental Note 4; Supplemental Fig. S11).

**Figure 6.**
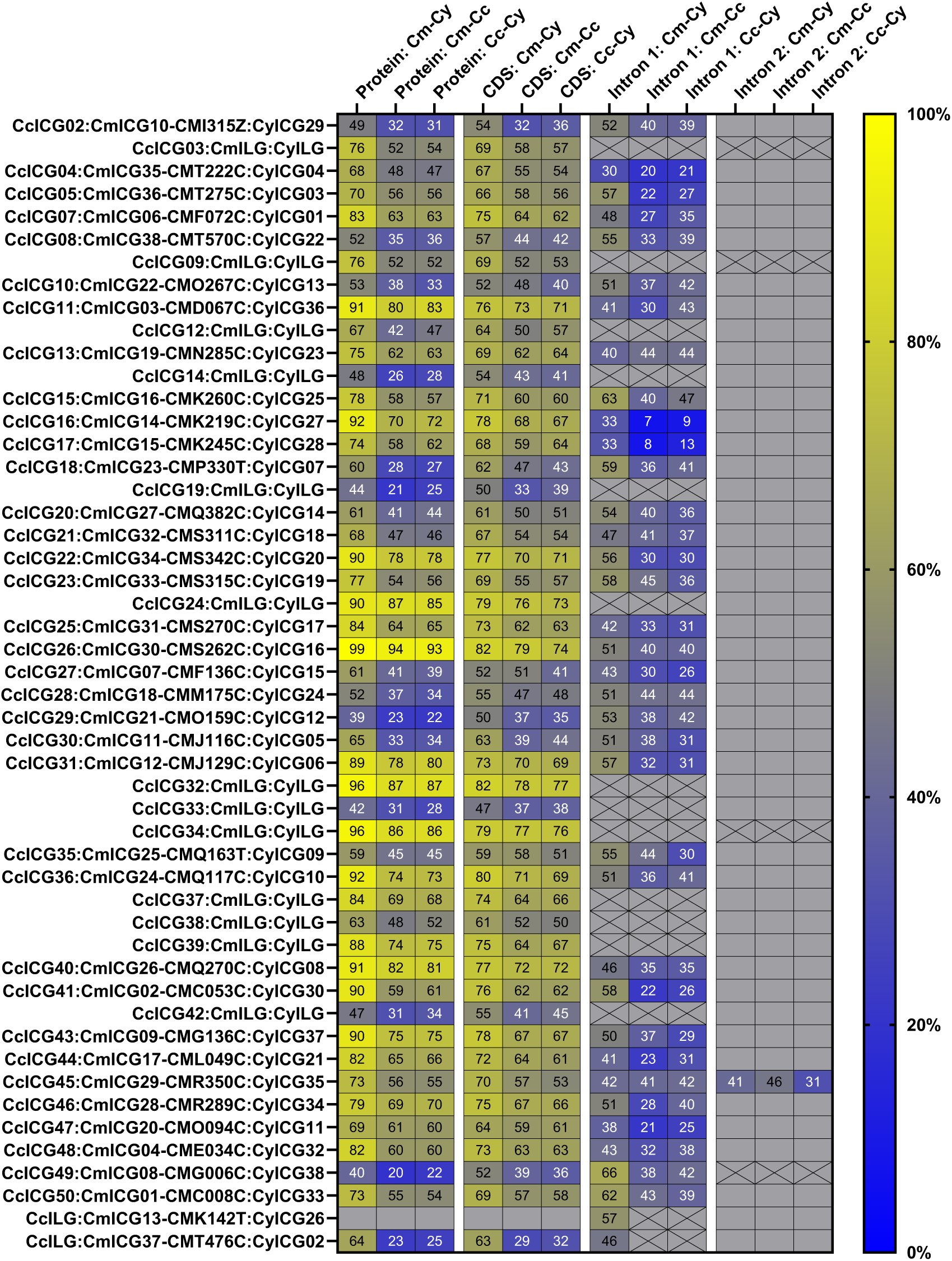
Heatmap of orthologous intron-containing genes in the Cx. Values indicate percent pairwise identity across the complete length of each protein, coding, or intron sequence. Intron-containing genes that are in-paralogs and intron-containing genes that lack orthologs have been omitted from the heatmap. Empty grey cells represent comparisons that are not applicable, while grey cells with a black X represent comparisons that are not possible due to intron loss. ICG, intron-containing gene; ILG, intron-lacking gene.

It has previously been observed that organisms with a reduced intron complement often have stronger conservation of splice sites. Analysis of sites among the Cx also revealed a highly constrained intron architecture. All species share a strongly conserved 5′ splice site core motif of ‘GUAAGU’, with species-specific extended 5′ motif variants, together with an almost invariant branch site consensus of ‘ACUAACC’ and a canonical, invariant 3′ splice site consensus of ‘AG’ (Supplemental Note 6; Supplemental Figs. S13–S16). Intron length and free energy were broadly similar among the species, whereas Cc introns had higher GC content, a longer branchpoint to 3′ splice site distance, and weaker 5′ splice sites (based on a scoring model trained on human sequences), while 3′ splice site scores did not differ significantly among the Cx (Supplemental Note 5; Supplemental Fig. S12).

To investigate whether introns have been retained in particular classes of gene, we performed functional classification of the ICGs across the Cx (Fig. 7). This revealed that 65% clustered within the following functional categories: translation, ribosomal structure, and ribosomal biogenesis (22 genes); post-translational modification, protein turnover, and chaperones (16 genes); energy production and conversion (13 genes); amino acid transport and metabolism (12 genes); as well as genes encoding proteins of unknown function (19 genes). The remaining 35% of ICGs were distributed over a range of functional categories. These results indicate that extreme intron depletion in the Cx has not occurred uniformly across the genome, but instead has preferentially retained introns within a restricted and functionally biased subset of genes.

**Figure 7.**
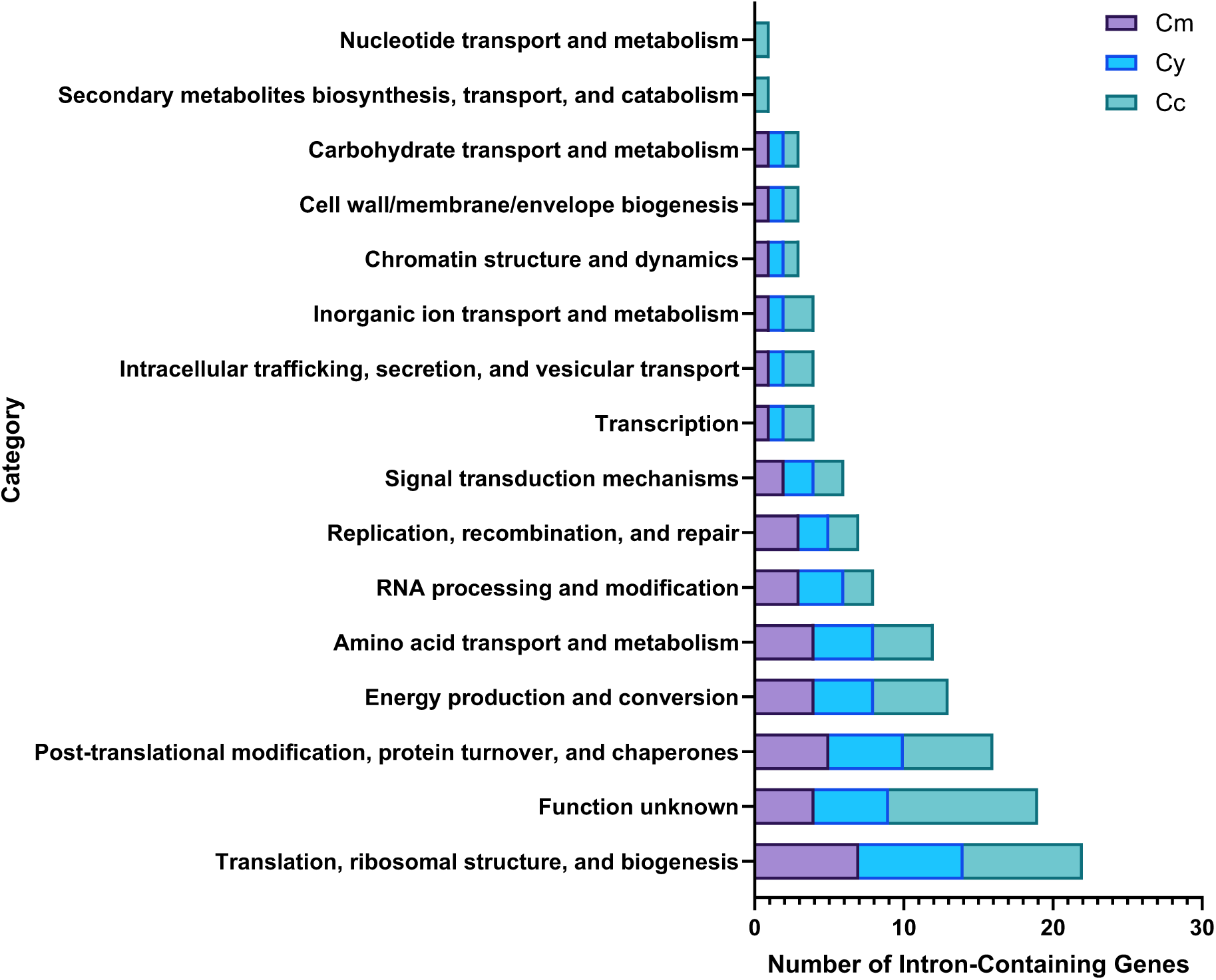
Functional classification of the Cx intron-containing genes. Intron-containing genes were classified according to the eukaryotic orthologous groups (KOG) categorization system.

### Splicing efficiency

The stark reduction in spliceosome complexity across the Cx raises the possibility of a corresponding decrease in splicing of intron-containing transcripts with possible consequences for the production of functional mRNA and proper gene expression. We therefore measured splicing efficiency, a measure of the extent to which introns are removed from pre-mRNA transcripts during the splicing process. Previous work established that pre-mRNA splicing in Cm is markedly inefficient relative to other eukaryotes (Schärfen et al. 2022; Wong et al. 2023). We investigated whether this pattern extends across the Cx by calculating the splicing efficiency of every Cx intron as a ratio ranging from 0 (no splicing) to 1 (complete splicing). Under standard laboratory growth conditions, splicing efficiency varied widely among the introns within each species, ranging from 0.03 to 0.87 (Supplemental Fig. S17A, B, C). Nonetheless, the average global splicing efficiencies are uniformly low (0.42, 0.50, and 0.42, respectively for Cm, Cy, Cc), with no statistically significant differences among the species (Fig. 8A). The distributions of splicing efficiencies are markedly bimodal in all three species. This result demonstrates that splicing inefficiency is shared among the Cx.

**Figure 8.**
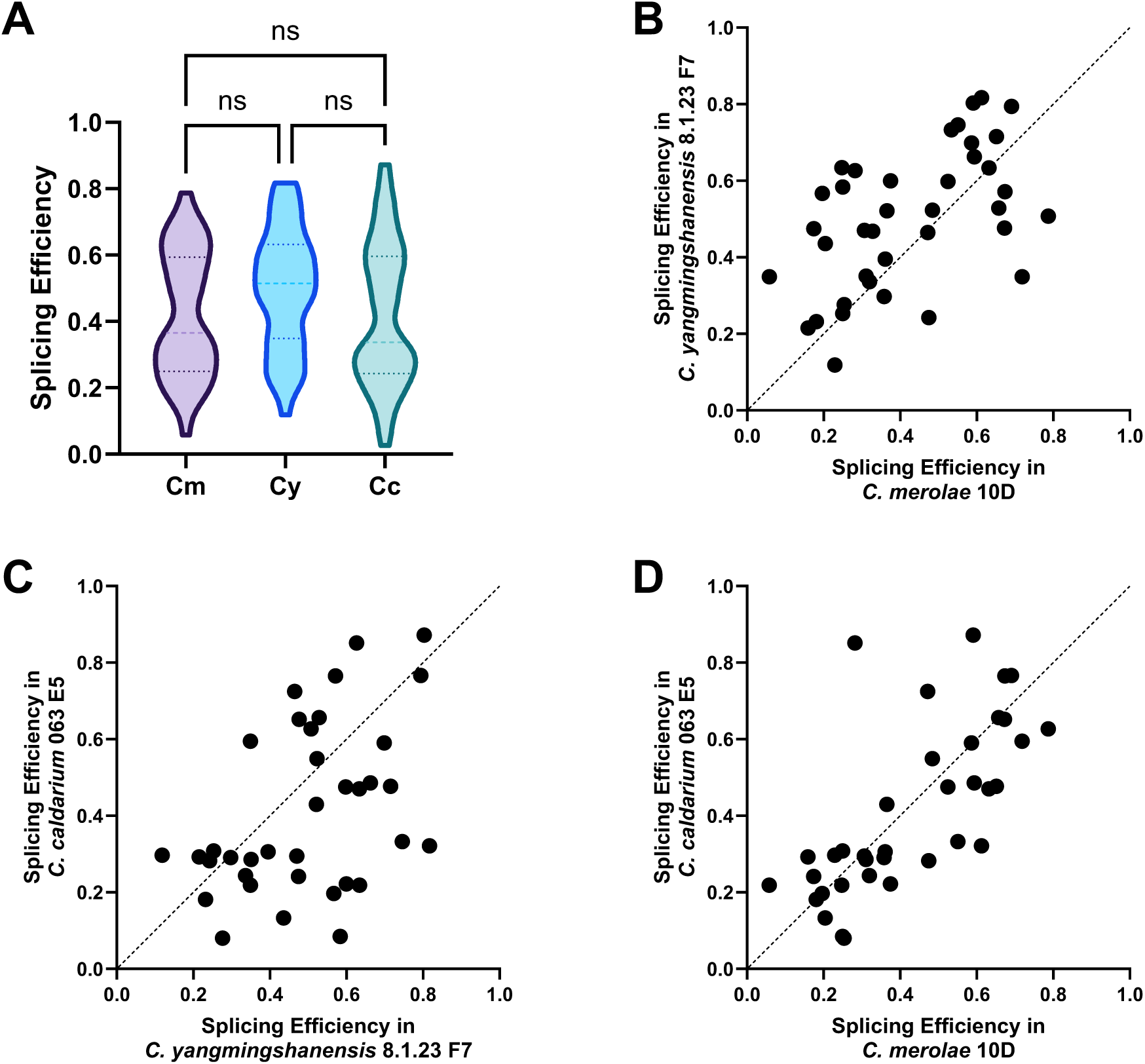
Comparison of splicing efficiency among the Cx under standard laboratory growth conditions. **(A)** Global splicing efficiency in *C. merolae* 10D (Cm), *C. yangmingshanensis* 8.1.23 F7 (Cy), and *C. caldarium* 063 E5 (Cc). An ordinary one-way ANOVA was conducted using Tukey’s multiple comparisons test, with a single pooled variance, to determine the significance of the multiple comparisons (ns = *p* > 0.05). **(B)** Each circle represents the splicing efficiency for one of the 38 introns that are comparable between Cm and Cy. Pearson correlation coefficient = 0.5435 (*p* = 0.0004). **(C)** Each circle represents the splicing efficiency for one of the 36 introns that are comparable between Cy and Cc. Pearson correlation coefficient = 0.4549 (*p* = 0.0053). Dashed line of identity is shown. **(D)** Each circle represents the splicing efficiency for one of the 36 introns that are comparable between Cm and Cc. Pearson correlation coefficient = 0.7044 (*p* < 0.0001).

If inefficient splicing is a stochastic event, a particular intron might be spliced well in one alga and poorly in another. Conversely, if splicing of individual introns is under selective pressure, a given intron would have similar levels of splicing across the Cx. To test this, we performed pairwise comparisons of orthologous intron splicing efficiencies. This revealed significant correlations between Cm and Cy (Fig. 8B; *r* = 0.54), and Cy and Cc (Fig. 8C; *r* = 0.45), with the strongest correlation occurring between Cm and Cc (Fig. 8D; *r* = 0.70). These findings suggest that despite potential species-specific adaptations, the splicing machinery and any intronic regulatory elements that influence splicing efficiency patterns have remained broadly conserved. Prompted by variation in splicing efficiency between introns within the same gene (Supplemental Fig. S17), we tested for correlations between splicing efficiency and intron features, but found no significant associations in Cm or Cy. In contrast, in Cc splicing efficiency was negatively correlated with intron length and positively correlated with intron free energy (Supplemental Note 7; Supplemental Figs. S18–S20).

Given the photosynthetic nature of the Cx, we tested the possibility that splicing levels are regulated by light by analyzing RNA-seq data from two previously published studies on Cm and one on Cy. In the Wong et al. (2023) Cm dataset, the average global splicing efficiencies were similar at 0.37 under dark and 0.32 under light, with no statistically significant difference observed on a global basis (Supplemental Note 8; Supplemental Fig. S21). In contrast, in the Abram et al. (2022) Cm dataset, average global splicing efficiencies were determined to be 0.32 under low light, 0.42 under moderate light, and 0.40 under extreme high light, with the global efficiency of splicing being significantly higher under moderate light than low light (*p* = 0.0199; Supplemental Note 9; Supplemental Fig. S22). In the Liu et al. (2020) Cy dataset, average global splicing efficiencies were found to be 0.26 for dark, 0.49 for low light, and 0.52 for high light, with global splicing efficiency significantly higher under both low and high light conditions than in the dark (*p* < 0.0001; Supplemental Note 10; Supplemental Fig. S23). These results point to modest changes in splicing in response to light in Cm, depending on the data used, with a more robust change in Cy.

In view of the variability in global changes in splicing to differing light levels, we tested whether individual introns are similarly spliced when we compared similar dark and light conditions. The strongest correlation between Cm and Cy was under dark conditions (Fig. 9A), where we found a moderate yet significant correlation between orthologous introns (*r* = 0.58, *p* = 0.0001), accompanied by a significantly higher global splicing efficiency in Cm (*p* = 0.0016). In low light conditions (Fig. 9B), the interspecies correlation weakened appreciably (*r* = 0.45, *p* = 0.0045), along with a global splicing efficiency that was markedly higher in Cy (*p* < 0.0001). Under high light exposure (Fig. 9C), an intermediate correlation strength was identified (*r* = 0.51, *p* = 0.0012), again coinciding with a significantly greater global splicing efficiency in Cy (*p* = 0.0054). From these results we conclude that orthologous introns are spliced at similar levels between Cm and Cy under all light conditions.

**Figure 9.**
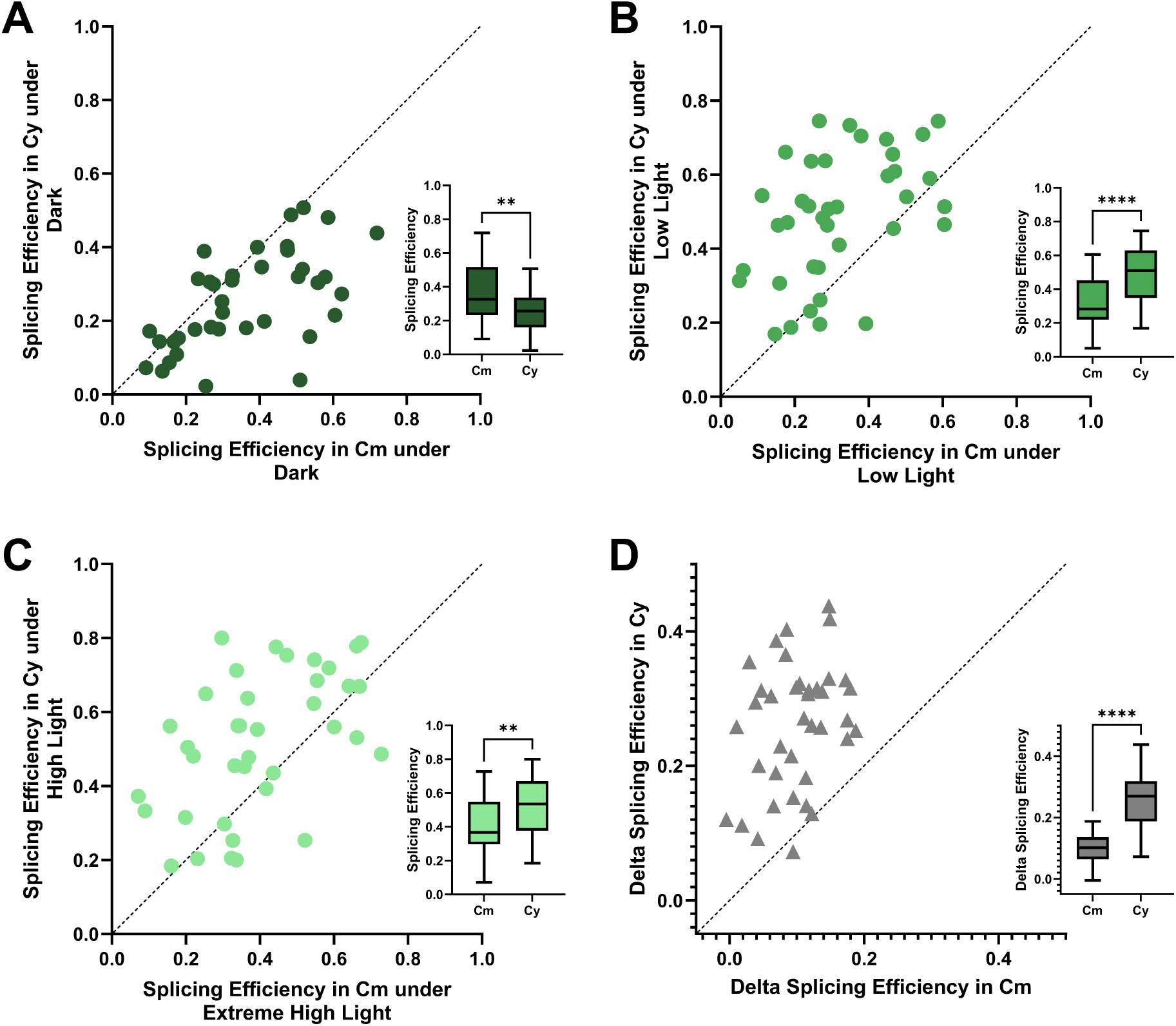
Interspecies comparisons of splicing efficiency between *C. merolae* 10D (Cm) and *C. yangmingshanensis* THAL066 (Cy) under dark, low light, and high light conditions. Dashed line of identity is shown. **(A)** Each circle represents the splicing efficiency for one of the 38 introns that are comparable between Cm and Cy under dark conditions. Pearson correlation coefficient = 0.5809 (*p* = 0.0001). The inset shows the global splicing efficiency of all 39 introns in Cm and all 40 introns in Cy under dark conditions. The significance of the difference was tested using an unpaired two-tailed *t*-test (** = *p* ≤ 0.01). **(B)** Each circle represents the splicing efficiency for one of the 38 introns that are comparable between Cm and Cy under low light conditions. Pearson correlation coefficient = 0.4509 (*p* = 0.0045). The inset shows the global splicing efficiency of all 39 introns in Cm and all 40 introns in Cy under low light conditions. The significance of the difference was tested using an unpaired two-tailed *t*-test (**** = *p* ≤ 0.0001). **(C)** Each circle represents the splicing efficiency for one of the 38 introns that are comparable between Cm and Cy under high light conditions. Pearson correlation coefficient = 0.5053 (*p* = 0.0012). The inset shows the global splicing efficiency of all 39 introns in Cm and all 40 introns in Cy under high light conditions. The significance of the difference was tested using an unpaired two-tailed *t*-test (** = *p* ≤ 0.01). **(D)** Comparison of the largest intraspecies light-induced splicing efficiency response between Cm and Cy. Each triangle represents the delta of the splicing efficiency for one of the 38 comparable introns going from low light to moderate light in Cm and from dark to high light in Cy. Pearson correlation coefficient = 0.3176 (*p* = 0.0520). The inset shows the delta splicing efficiency values for Cm and Cy. The significance of the difference was tested using an unpaired two-tailed *t*-test (**** = *p* ≤ 0.0001).

To test whether the response of some introns to changes in light is more highly conserved than the absolute level, we compared the delta of the largest intraspecies light-induced splicing efficiency responses between Cm (low to moderate light) and Cy (dark to high light). This revealed only a marginal correlation among the orthologous introns (Fig. 9D; *r* = 0.32, *p* = 0.05). On a global basis, however, the magnitude of the splicing response was significantly higher in Cy (*p* < 0.0001). Together, these results indicate that although orthologous introns in Cm and Cy retain broadly similar splicing efficiency patterns across matched conditions, the magnitude of light-responsive splicing differs between species and is more pronounced in Cy.

One possible explanation of the low splicing levels across the Cx is that there is extensive alternative splicing. We inspected the Sashimi plots of all ICGs in the Cx for evidence of alternative splicing in Cm, Cy, and Cc (Supplemental Figs. S24–S27) over all conditions. Intron retention is the predominant form of alternative splicing in the Cx, with all introns exhibiting retention to varying degrees, which is congruent with the splicing efficiencies observed. Intriguingly, 15 ICGs in Cm, 13 in Cy, and 9 in Cc displayed pronounced intron accumulation, for which intronic read coverage exceeds that of the surrounding exons. Additional forms of alternative splicing are detectable but rare across all three species. Specifically, alternative 3′ splice site usage was observed in three ICGs in Cm, four in Cy, and three in Cc; exon skipping occurred in one ICG in Cm, two in Cy, and three in Cc; and alternative 5′ splice site usage was demonstrated in one ICG in Cc.

Finally, we sought to explain how the Cx cope with such low splicing efficiencies. One possibility is simply that their ICGs are up-regulated to compensate for the unspliced fraction. Analysis of transcriptomic data showed that ICGs were significantly more highly expressed than intron-lacking genes (ILGs) in all three species (Fig. 10A, C, E). Median log₂(TPM + 1) values for ICGs and ILGs were 8.2 and 6.2 in Cm, 8.1 and 5.8 in Cy, and 7.9 and 6.5 in Cc, respectively (*p* < 0.0001 for all comparisons). After adjusting ICG TPM values by gene-specific splicing efficiency (SE), the difference between ICGs and ILGs was reduced, but remained significant in each species (Fig. 10B, D, F). Median adjusted expression values, calculated as log₂(TPM × SE + 1) for ICGs, were 6.6 and 6.2 in Cm (*p* = 0.01), 7.0 and 5.8 in Cy (*p* = 0.0002), and 6.9 and 6.5 in Cc (*p* = 0.03).

**Figure 10.**
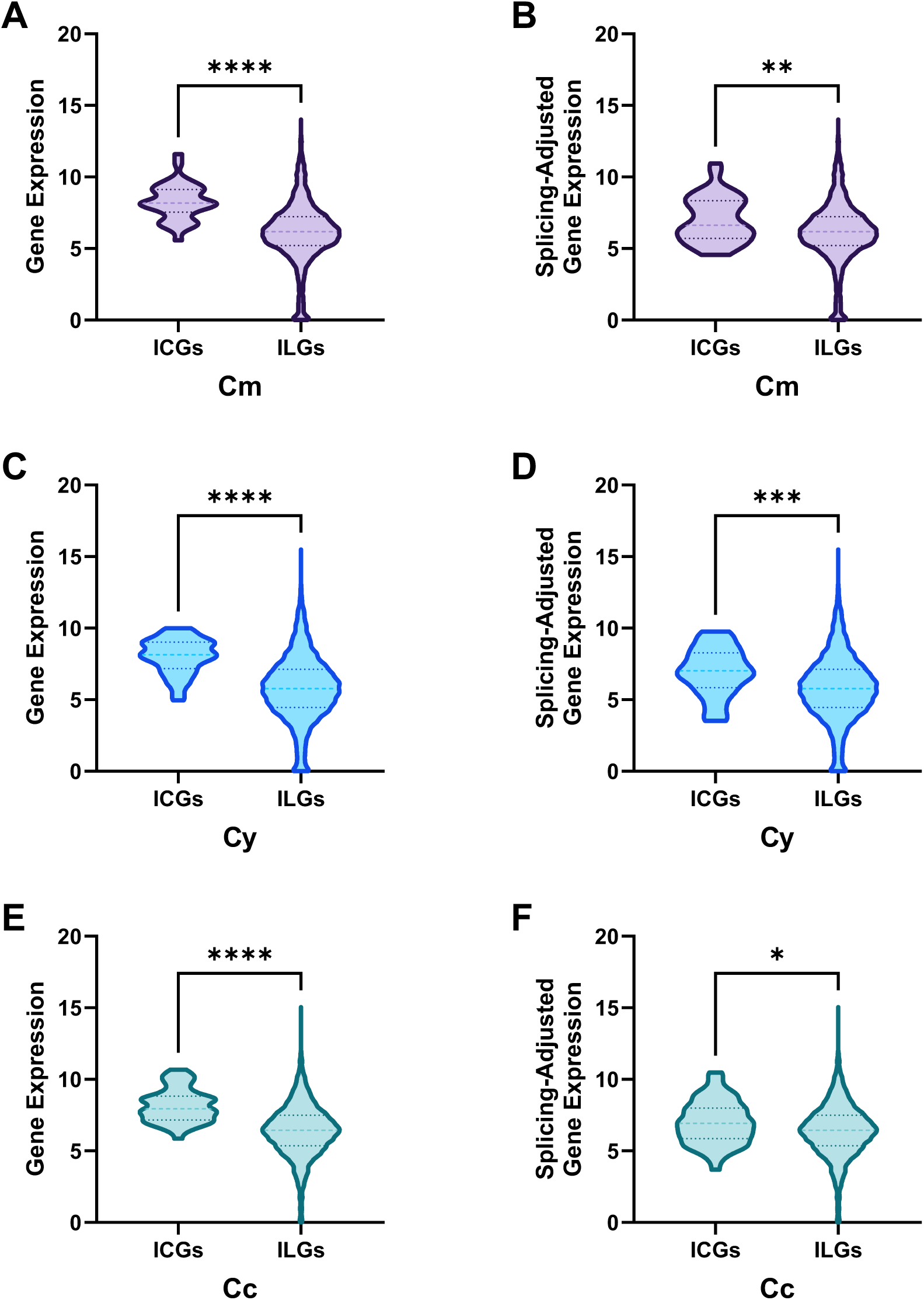
Expression distributions of intron-containing and intron-lacking genes in the Cx. Gene-level expression of intron-containing genes (ICGs) and intron-lacking genes (ILGs) in *C. merolae* 10D (Cm), *C. yangmingshanensis* 8.1.23 F7 (Cy), and *C. caldarium* 063 E5 (Cc). Violin plots depict Cm gene expression **(A)** and splicing-adjusted gene expression **(B)**, Cy gene expression **(C)** and splicing-adjusted gene expression **(D)**, and Cc gene expression **(E)** and splicing-adjusted gene expression **(F)**. For ICGs, gene expression is shown as log₂(TPM + 1), and splicing-adjusted gene expression is shown as log₂(TPM × SE + 1), where SE denotes gene-specific splicing efficiency. For ILGs, values are shown as log₂(TPM + 1) in all plots. Statistical comparisons were performed using two-tailed Mann–Whitney tests; * = *p* ≤ 0.05, ** = *p* ≤ 0.01, *** = *p* ≤ 0.001, and **** = *p* ≤ 0.0001.

## Discussion

We previously reported that Cm possesses a drastically reduced spliceosome, an extremely small intron repertoire, and unusually low splicing efficiency (Stark et al. 2015; Reimer et al. 2017; Schärfen et al. 2022; Black et al. 2023; Wong et al. 2023). By extending this analysis to Cy and Cc, we demonstrate that these features are not unique to Cm, but instead, define the Cx as a whole.

### Genome streamlining and GC bias

Compared to the last common ancestor of the Rhodophyta, Cm is inferred to have undergone a net loss of more than 1,700 orthologous gene families, suggesting that its compact nuclear genome of 17 Mbp is the result of substantial reductive evolution (Qiu et al. 2013; Miyagishima and Tanaka 2021). Although Cy and Cc encode a similar number of genes as Cm, their genomes are even more compact, with Cy approximately three quarters the size of Cm and Cc roughly half the size of Cm, explained by differences in subtelomeric duplication, repeat expansion, and reduced intergenic DNA among the Cx (Matsuzaki et al. 2004; Cho et al. 2023). Despite the genome size difference between Cm and Cy, both species share a similar GC content of 55%, whereas Cc has a much higher GC content of 66%. This compositional bias permeates its splicing landscape, influencing amino acid usage in splicing proteins, snRNA stability, coding sequence composition, intron GC content, and splice site architecture (Supplemental Note 11). Thus, genome composition directly shapes spliceosomal and intronic features within the Cx.

### A shared, highly reduced spliceosome

All splicing proteins present in Cm have orthologs in Cy and Cc, and only a small subset shows above-average conservation across species, typically those with additional known RNA-processing functions. Most importantly, U1 snRNA and its associated proteins are absent from all three species. This establishes U1 loss as a lineage-defining feature and indicates that extreme spliceosome reduction predates Cx divergence about 320 million years ago.

We detected only a small number of additional splicing protein candidates in Cy and Cc. This indicates that any potential lineage-specific retention of known splicing factors is modest and that the overall Cx splicing protein repertoire remains highly reduced. The 65 broadly conserved splicing proteins shared among the Cx species are far fewer than the 145 spliceosomal proteins inferred to have been present in the last eukaryotic common ancestor (Vosseberg et al. 2023).

Despite U1 loss, core snRNAs retain conserved catalytic motifs, indicating preservation of the essential spliceosome framework. The apparent absence of LSm8 across the Cx further supports reliance on a simplified LSm complex functioning in both splicing and mRNA turnover (Supplemental Note 12). Together, these findings define a stable, streamlined spliceosome operating without U1.

### Extreme intron depletion with constrained architecture

The Cx genomes contain only 39–54 introns, placing them among the most intron-poor free-living eukaryotes. The paucity of introns in the Cx represents a stark contrast to intron-rich species such as *H. sapiens* with 207,344 introns (Sakharkar et al. 2004), the fruit fly *Drosophila melanogaster* with 48,257 introns (Misra et al. 2002), the extremophilic red algae *G. yellowstonensis* (Cho et al. 2023) and *G. sulphuraria* (Schönknecht et al. 2013) with 15,190 and 13,630 introns, respectively, as well as the fission yeast *Schizosaccharomyces pombe* with 4,730 introns (Wood et al. 2002). The Cx have a markedly lower number of introns, even compared to other intron-poor species such as the budding yeast *S. cerevisiae* with 296 introns (Parenteau et al. 2008) and the mesophilic red alga *Porphyridium purpureum* with 235 introns (Bhattacharya et al. 2013), suggesting an exceptional level of genome streamlining. Indeed, to find species with fewer introns than members of the Cx, one must look beyond free-living eukaryotes to parasitic or endosymbiotic lineages such as the parasitic microsporidian *Encephalitozoon cuniculi*, which possesses only 36 introns (Lee et al. 2010), the nucleomorph of the cryptomonad alga *Guillardia theta*, with just 17 introns (Douglas et al. 2001), and the parasitic diplomonad *Giardia intestinalis*, with a mere 13 introns (Xu et al. 2020). The paucity of Cx introns may not be surprising given the highly reduced nature of the Cx spliceosome, consistent with the finding that the intron-poor yeast *S. cerevisiae* has lost more than 40 spliceosomal proteins that are largely maintained in other intron-rich fungal lineages (Sales-Lee et al. 2021).

This minimal intron complement in the Cx is accompanied by highly conserved splicing motifs similar to those found in hemiascomycetous yeasts such as *S. cerevisiae* (Irimia and Roy 2008). We found that the highly conserved ‘GUAAGU’ consensus sequence of the Cx 5′ splice sites is consistent with most intron-poor eukaryotes and reflects efficient selection for this strong consensus sequence (Irimia et al. 2009). This is notable in light of the absence of the U1 snRNA in this lineage: the 5′ splice site conservation does not simply reflect strong binding to U1. We further observed that the Cx 3′ splice site sequence is an invariant and canonical ‘AG’, while the branch site is a virtually invariant ‘ACUAACC’. Other intron-poor species also possess similarly strict branch site sequences and tend to exhibit highly regular intronic sequences (Irimia and Roy 2008). Such homogenization of the Cx splicing signals provides further support for a positive correlation between spliceosome complexity and intron diversity (Sales-Lee et al. 2021), and is consistent with co-evolution of intron simplification and spliceosome reduction.

Although overall intron sequence conservation is lower than that of coding regions, certain introns exhibit unexpectedly high conservation, indicating locus-specific constraint despite global intron loss. Moreover, we found a prominent 5′ bias for PTCs across the Cx. Notably, phase 2 introns were the most prevalent across the Cx, and since they interrupt the coding sequence after the second nucleotide of a codon, those conforming to the 5′ splice site consensus sequence of ‘GUAAGU’ would possess an in-frame ‘UAA’ termination codon when retained. Although many introns harbour in-frame PTCs that can target intron-retaining transcripts for nonsense-mediated decay (NMD) (Monteuuis et al. 2019), it is possible that Cm lacks a functional NMD pathway due to absent or heavily derived NMD factors (Lloyd and Davies 2013; Lloyd 2018). In this scenario, the frequent occurrence of early intronic PTCs in the Cx may serve as an additional quality-control safeguard against aberrant protein production from unspliced transcripts in a system characterized by inefficient splicing.

The large number of ICGs found to be involved in translation, ribosomal structure, and ribosomal biogenesis is noteworthy, as these genes are central to fundamental cellular processes and may benefit from hosting introns, which can enhance transcriptional and translational yield, as shown in *S. cerevisiae* (Juneau et al. 2006). Moreover, the abundance of genes with unknown functions highlights the complexity and relatively understudied nature of these extremophilic algae, suggesting possible novel roles for genes awaiting functional characterization.

### Conserved splicing inefficiency modulated by light

Global splicing efficiency is uniformly low across the Cx, with average values of 0.42, 0.50, and 0.42 in Cm, Cy, and Cc, respectively. This relatively low level of splicing efficiency stands in sharp contrast to other species with highly efficient splicing, such as *H. sapiens* (Tilgner et al. 2012), the fruit fly *D. melanogaster* (Khodor et al. 2011), the budding yeast *S. cerevisiae* (Oesterreich et al. 2016), and the fellow red alga *G. sulphuraria* (Wong et al. 2023). Notably, using a method virtually identical to the one employed in this study, the average global splicing efficiency for the fission yeast *S. pombe* was found to be 0.90 (Porat et al. 2023). Moreover, the splicing efficiencies of orthologous introns are significantly correlated between species, indicating that intron-specific determinants of inefficient splicing have been maintained over evolutionary time. The reduced splicing efficiency observed in the Cx is consistent with substantial loss of spliceosomal components and may reflect limitations in overall splicing capacity and, potentially, splicing accuracy (Sales-Lee et al. 2021; Black et al. 2023).

Light modulates splicing efficiency in Cm and Cy, with stronger responses in Cy, demonstrating that the reduced spliceosome remains environmentally responsive. This response is substantial, with global splicing efficiency increasing by up to 31% in Cm and 100% in Cy. While there is no evidence to suggest that Cc would deviate substantially from these patterns, given the overall similarities in the Cx splicing landscape, the absence of the transcriptomic data required to make a definitive determination for Cc is nonetheless a limitation of this study. Intron retention is the predominant alternative splicing outcome across the Cx, consistent with globally inefficient intron removal. Intron retention is also the most prevalent form of alternative splicing in the rice *Oryza sativa* (Wang and Brendel 2006), the thale cress *Arabidopsis thaliana* (Ner-Gaon et al. 2004), and the unicellular marine diatom *Phaeodactylum tricornutum* (Rastogi et al. 2018).

Collectively, our results indicate that extensive intron loss, spliceosomal protein reduction, and elimination of the U1 snRNA occurred prior to the divergence of the Cx approximately 320 million years ago. The complex interplay of reductive co-evolution between the Cx spliceosome and its intron substrates has produced a streamlined system with a minimal catalytic core, highly constrained intron architecture, and relatively inefficient splicing that nevertheless remains viable and environmentally responsive. These findings support the idea that once intron number is drastically reduced and splice site signals become highly homogenized, selective pressure to maintain a fully elaborated spliceosome may be relaxed, permitting the evolutionary loss of spliceosomal components traditionally considered essential, and allowing for the stable persistence of a simplified yet functional system. Rather than representing degeneration, the Cx spliceosome appears to constitute a stable evolutionary configuration compatible with extreme intron depletion. In this sense, the Cx provide a natural evolutionary experiment demonstrating that the spliceosome is not a fixed entity, but a modular system whose complexity can scale with intron burden. These extremophilic red algae may therefore define a lower bound for spliceosomal complexity in free-living eukaryotes and provide a framework for understanding how core RNA-processing machineries can be extensively streamlined while remaining functional.

## Materials and Methods

### Data and quality assessment

The *Cyanidioschyzon merolae* 10D reference genome (Matsuzaki et al. 2004; Nozaki et al. 2007) was obtained from release 56 of Ensembl Plants (Yates et al. 2022) and NCBI GenBank (accession numbers GCA_000091205.1, AB002583.1, and D89861.1 for the nuclear, chloroplast, and mitochondrion genomes, respectively). The *Cyanidiococcus yangmingshanensis* THAL066 reference genome (Liu et al. 2020) was obtained from NCBI GenBank (accession numbers GCA_013995675.1, MN431657.1, and MN431656.1 for the nuclear, chloroplast, and mitochondrion genomes, respectively). The nuclear reference genomes (Cho et al. 2023) of the following species were downloaded from NCBI GenBank: *Cyanidium caldarium* 063 E5 (GCA_026184775.1), *Cyanidiococcus yangmingshanensis* 8.1.23 F7 (GCA_026122185.1), and *Galdieria yellowstonensis* 108.79 E11 (GCA_026122205.1).

Supplemental Table S10 lists the RNA sequencing (RNA-seq) data used in this study that were downloaded from the National Center for Biotechnology Information (NCBI) Sequence Read Archive (SRA) (Sayers et al. 2021).

The raw Illumina RNA-seq reads in FASTQ format were evaluated for quality using FastQC v0.12.1 (https://www.bioinformatics.babraham.ac.uk/projects/fastqc) and MultiQC v1.21 (Ewels et al. 2016).

### Identification of splicing proteins

To identify the complement of splicing proteins present in *C. caldarium* 063 E5 and *C. yangmingshanensis* 8.1.23 F7, a reciprocal best hits strategy was employed as a proxy for orthology (Ward and Moreno-Hagelsieb 2014) using BLAST+ v2.14.1 (Camacho et al. 2009). Forward searches were conducted using the tblastn application with *C. merolae* 10D splicing protein sequences identified previously (Stark et al. 2015; Reimer et al. 2017; Black et al. 2023) queried against a custom BLAST subject database consisting of either the complete *C. caldarium* 063 E5 or *C. yangmingshanensis* 8.1.23 F7 reference genome. The corresponding reverse searches were conducted using the blastp application with the protein sequence hits queried against *C. merolae* 10D entries in the nr database (October 18, 2023) (Sayers et al. 2021).

In order to rule out the presence of any splicing proteins in *C. caldarium* 063 E5 or *C. yangmingshanensis* 8.1.23 F7 that may be absent in *C. merolae* 10D, an exhaustive list of splicing proteins from *Arabidopsis thaliana* (Taxonomy ID 3702), *Homo sapiens* (Taxonomy ID 9606), *Saccharomyces cerevisiae* (Taxonomy ID 559292; strain ATCC 204508 / S288C), and *Schizosaccharomyces pombe* (Taxonomy ID 284812; strain 972h- / ATCC 24843) was compiled. The QuickGO browser (Binns et al. 2009) was used to obtain the reviewed Swiss-Prot entries from the UniProt Knowledgebase (The UniProt Consortium 2020) that were associated by the Gene Ontology Annotation (GOA) project (Huntley et al. 2014) with the Gene Ontology (GO) terms (The Gene Ontology Consortium 2020) “mRNA splicing, via spliceosome” (GO:0000398), “spliceosomal complex” (GO:0005681), “pre-mRNA 5′-splice site binding” (GO:0030627), “pre-mRNA 3′-splice site binding” (GO:0030628), “RES complex” (GO:0070274), “nuclear mRNA surveillance of spliceosomal pre-mRNA splicing” (GO:0071030), “spliceosomal snRNP complex” (GO:0097525), and “splicing factor binding” (GO:1990935) (GO version 2022-04-24; annotation set created on 2022-03-24 11:08). Forward searches were conducted using the tblastn application with the Swiss-Prot splicing protein sequences being queried against the custom BLAST subject *C. caldarium* 063 E5 or *C. yangmingshanensis* 8.1.23 F7 database, while the reverse searches were conducted using the blastp application with the initial hits being queried against the respective species entries in the swissprot database (October 18, 2023) (Sayers et al. 2021). Additional splicing protein candidates were required to lack a reciprocal best hit in *C. merolae* 10D, to have a reciprocal best hit in *C. caldarium* 063 E5 or *C. yangmingshanensis* 8.1.23 F7, and to meet a maximum E-value threshold of 1 × 10⁻⁵ for inclusion.

### Identification of snRNAs

The covariance models for the U1, U2, U4, U5, and U6 snRNAs (accession numbers RF00003, RF00004, RF00015, RF00020, and RF00026, respectively) were retrieved from release 14.10 of the Rfam database (Kalvari et al. 2020). The cmalign program of Infernal v1.1.5 (Nawrocki and Eddy 2013) was used to align the U2, U4, U5, and U6 snRNA sequences from *C. merolae* 10D (Stark et al. 2015) to each respective covariance model without the use of HMM banding (--nonbanded), followed by the building (cmbuild) and calibrating (cmcalibrate) of the augmented covariance models. The cmsearch program was then used to search each covariance model against the Cyanidiales and Cyanidioschyzonales genomes with options to maximize sensitivity and encourage more full-length hits (--max --nohmmonly) in order to identify snRNA orthologs. The top Infernal hit for the U1 snRNA covariance model in *C. merolae* 10D (E-value of 0.59 and score of 16.1) was used as the inclusion threshold for the other Cyanidiales and Cyanidioschyzonales species. The top hit in *C. yangmingshanensis* 8.1.23 F7 (E-value of 1.1 and score of 14.8) and the top hit in *C. caldarium* 063 E5 (E-value of 1.8 and score of 13.5) failed to meet this threshold.

Secondary structure models of the snRNAs for *C. caldarium* 063 E5 and *C. yangmingshanensis* 8.1.23 F7 were generated manually using RNAfold v2.6.3 (Lorenz et al. 2011) for rapid identification of potential base-paired regions and Adobe Illustrator v29.2.1 (https://www.adobe.com) for model-building to ensure consistency with structures in the literature, as described previously (Stark et al. 2015). Compared to the Infernal hits for *C. yangmingshanensis* 8.1.23 F7 snRNAs, the 3′ end of U4 was shortened by 1 nt, and the 5′ and 3′ ends of U5 were extended by 132 and 238 nt, respectively. For *C. caldarium* 063 E5 snRNAs, the 3′ end of U4 was extended by 226 nt, and the 5′ and 3′ ends of U5 were extended by 224 and 186 nt, respectively.

### Identification of intron-containing genes

The raw Illumina RNA-seq reads from all samples cultured under standard laboratory growth conditions (Supplemental Table S10) were aligned using two different aligners, HISAT2 v2.2.1 (Kim et al. 2019) and STAR v2.7.10b (Dobin et al. 2012). The indices for HISAT2 and STAR were built without the use of annotations.

For the HISAT2 alignments, the minimum allowed intron length was set to 20 bp (--min-intronlen 20) and the maximum allowed intron length was set to 10,000 bp (--max-intronlen 10000). In addition, the input of strand-specific, paired-end reads was indicated (--rna-strandness RF) and the downstream transcriptome assembly option (--dta) was used in order to decrease the number of false positive splice junctions by requiring longer anchor lengths for splice sites.

Since HISAT2 outputs alignments in the SAM format, it was necessary to convert the alignments to the BAM format with SAMtools v1.17 (Danecek et al. 2021) for downstream analysis.

For the STAR alignments, the minimum allowed intron size was set to 20 bp (--alignIntronMin 20), the maximum allowed intron size was set to 10,000 bp (--alignIntronMax 10000), and the maximum allowed gap between two mates was set to 10,000 bp (--alignMatesGapMax 10000). Furthermore, the two-pass mapping mode was utilized (--twopassMode Basic) in order to increase the robustness of splice junction discovery. The alignments were output as unsorted BAM files (--outSAMtype BAM Unsorted).

Portcullis v1.2.4 (Mapleson et al. 2018) was used to compile a high-confidence set of splice junctions from the HISAT2 and STAR alignments. The Portcullis sub-tool, full, was used to execute all three stages of the Portcullis pipeline (namely, prepare, junction analysis, and junction filtering). Portcullis was run using CSI indexing (--use_csi), the orientation and strandedness of the reads that produced the BAM alignments were specified (--orientation FR --strandedness firststrand), the option to calculate additional metrics was used (--extra), and only junctions with a number of split reads greater than or equal to 2 were kept (--min_cov 2).

The sequences of the potential introns, extracted from each reference genome using BEDTools v2.31.1 (Quinlan and Hall 2010), were examined for the presence of the putative branchpoint consensus sequence ‘ACTAACC’ (Matsuzaki et al. 2004), as well as to determine the 5′ and 3′ splice site sequences. The lowest value for the score and the lowest value for the number of spliced alignments among the introns possessing the putative branchpoint consensus sequence was used to establish a threshold whereby any potential intron with lower values was discarded. Stringency was further increased by requiring potential introns to be present in both the HISAT2 and STAR alignments for each RNA-seq sample. All remaining potential introns were then subjected to visual inspection.

The 5′ splice sites (3 bases in the exon and 6 bases in the intron) and the 3′ splice sites (20 bases in the intron and 3 bases in the exon) were scored with MaxEntScan (Yeo and Burge 2004) using a maximum entropy model. Sequence logos were generated using Geneious Prime v2025.0.3 (https://www.geneious.com) and WebLogo v3.7.12 (Crooks et al. 2004).

Intron-containing genes were classified according to the eukaryotic orthologous groups (KOG) categorization system (Tatusov et al. 2003), using the highest-scoring eukaryotic functional category assigned by eggNOG v5.0 (Huerta-Cepas et al. 2019).

### Splicing efficiency

The intron retention ratios were calculated for all Illumina RNA-seq samples (Supplemental Table S10) using IRFinder-S v2.0.1 (Lorenzi et al. 2021). Differential intron retention between conditions was analyzed using the DESeq2 option (Love et al. 2014). The intron retention ratio for each intron was subtracted from 1 to calculate the splicing efficiency of each intron. Sashimi plots (Katz et al. 2015) were produced with the ggsashimi v1.1.5 tool (Garrido-Martín et al. 2018) using the IRFinder-S output.

### Gene expression

To evaluate gene-level expression differences between all intron-containing nuclear protein-coding genes and annotated intron-lacking nuclear protein-coding genes across *C. merolae* 10D, *C. yangmingshanensis* 8.1.23 F7, and *C. caldarium* 063 E5, STAR alignments from all samples cultured under standard laboratory growth conditions (Supplemental Table S10) were used for gene-level read counting with featureCounts v2.0.6 (Liao et al. 2013). Paired-end read (-p) fragments were counted (--countReadPairs) in reversely stranded mode (-s 2), requiring both ends to be aligned (-B) and excluding chimeric fragments (-C). Multi-mapping and multi-overlapping fragments were not counted. Feature lengths reported in the count output were used for TPM normalization (Wagner et al. 2012). Gene-level counts were converted to gene-level transcripts per million (TPM) values independently for each RNA-seq library. For each gene, fragment counts were divided by the corresponding feature length in kilobases. These length-normalized values were then divided by the sum of all length-normalized values within each library and multiplied by 1,000,000. For *C. merolae* 10D, which had three libraries, a representative TPM value for each gene was calculated as the mean TPM across libraries. For *C. yangmingshanensis* 8.1.23 F7 and *C. caldarium* 063 E5, each of which had one library, the representative TPM value corresponded to the single available library. The expression analysis included 4,806 genes for *C. merolae* 10D (37 intron-containing genes [ICGs] and 4,769 intron-lacking genes [ILGs]), 4,833 genes for *C. yangmingshanensis* 8.1.23 F7 (37 ICGs and 4,796 ILGs), and 4,871 genes for *C. caldarium* 063 E5 (50 ICGs and 4,821 ILGs). Gene expression values were transformed as log₂(TPM + 1) prior to visualization and statistical analysis. For splicing-adjusted gene expression, ICG TPM values were multiplied by gene-specific splicing efficiency (SE) prior to log transformation, yielding log₂(TPM × SE + 1). ILG values were not adjusted and remained log₂(TPM + 1).

### Determination of orthology, synteny, and average amino acid identity

In addition to relying on multiple reciprocal BLAST searches as a proxy for orthology, conducted using BLAST+ v2.12.0 custom databases as implemented within Geneious Prime, likely orthologs were further confirmed by synteny. To this end, SyMAP v5.4.9 (Soderlund et al. 2011), which incorporates the MUMmer program (Marçais et al. 2018), was used to compute and visualize synteny among the nuclear chromosomes.

The sequences for all annotated proteins in the genomes of *C. merolae* 10D, *C. caldarium* 063 E5, *C. yangmingshanensis* 8.1.23 F7, and *G. yellowstonensis* 108.79 E11 were used to determine the average amino acid identity (AAI) among these species using the EzAAI v1.2.3 (Kim et al. 2021) pipeline. The AAI values were calculated using the MMSeqs2 program (Steinegger and Söding 2017) with minimal constraints for identity and query coverage (-p mmseqs -id 0.01 -cov 0.01), followed by hierarchical clustering of taxa to create a phylogenetic tree that was visualized with Geneious Prime.

### Determination of transcriptional boundaries

To ascertain the transcriptional boundaries for genes of interest, the PacBio RNA-seq samples (Supplemental Table S10) for *C. merolae* 10D (Schärfen et al. 2022) were processed using PacBio’s IsoSeq v3.7.0 pipeline (https://github.com/PacificBiosciences/IsoSeq). The refine function of IsoSeq was used for the trimming of poly(A) tails (--require-polya) and the removal of concatemers based on the 5′ primer (GCAATGAAGTCGCAGGGTTGGG) and 3′ primer (GTACTCTGCGTTGATACCACTGCTT) used. The cluster function was then applied for iterative cluster merging and polished sequence generation using a consensus approach guided by quality values (--use-qvs). Polished isoforms with a predicted accuracy ≥ 0.99 were aligned to the *C. merolae* 10D reference genome using minimap2 v2.24 (Li 2021), with options selected to output alignments in the SAM format (-a), to use the preset for long-read splice alignment of PacBio IsoSeq or traditional cDNA reads (-x splice:hq), and to find canonical splicing sites on the transcript strand (-uf). PacBio RNA-seq samples belonging to the other species did not require processing with IsoSeq and proceeded directly to alignment with minimap2. The SAM output files were converted to coordinate-sorted and indexed BAM files using SAMtools.

In order to accurately determine the transcriptional boundaries for genes of interest that have low coverage depth in the PacBio RNA-seq samples, Trinity v2.14.0 (Haas et al. 2013) was used to perform both genome-guided and genome-free *de novo* transcriptome assembly. For the genome-guided option, the STAR output produced earlier for the Portcullis pipeline was coordinate-sorted by SAMtools and provided to Trinity for a genome-guided *de novo* transcriptome assembly (--genome_guided_bam). The maximum allowed intron size was set to 10,000 bp (--genome_guided_max_intron 10000), the strand-specific RNA-seq read orientation was specified (--SS_lib_type RF), the option to minimize the number of falsely fused transcripts when working with a gene-dense compact genome was exercised (--jaccard_clip), and the *in silico* normalization of reads was not performed (--no_normalize_reads) since computational considerations were not a factor. For the genome-free option, the raw Illumina RNA-seq reads from all standard samples (Supplemental Table S10) were subjected to adapter clipping as well as quality trimming by Trimmomatic v0.39 (Bolger et al. 2014) using settings (“ILLUMINACLIP:TruSeq3-PE.fa:2:30:10 SLIDINGWINDOW:4:5 LEADING:5 TRAILING:5 MINLEN:25”) recommended by Haas et al. (2013) and based on the work performed by MacManes (2014). Reads that were still paired after processing were provided to Trinity for a genome-free *de novo* transcriptome assembly (--seqType fq). The remaining applicable Trinity options were set as before (--SS_lib_type RF --jaccard_clip --no_normalize_reads). All Trinity output was aligned to the appropriate reference genome with minimap2, as described earlier.

### Visualization, alignments, and properties

Data were visualized using IGV v2.17.4 (Thorvaldsdóttir et al. 2012) and Geneious Prime. Transcriptional boundaries were used for the purpose of verifying, revising, or annotating coding and noncoding sequences.

Protein sequences were aligned using Clustal Omega v1.2.2 (Sievers and Higgins 2018), as implemented in Geneious Prime, with options set automatically (--auto) and the sequence type to protein (-t Protein). Nucleic acid alignments were generated using the iterative refinement algorithm G-INS-i (--globalpair) of MAFFT v7.490 (Katoh and Standley 2013), as implemented by the Geneious Prime plugin.

The free energy of the thermodynamic ensemble for each snRNA and intron was computed using RNAfold v2.6.4 (Lorenz et al. 2011). Unless otherwise indicated, Geneious Prime was used to calculate all nucleic acid and amino acid sequence properties.

### Statistical analysis

Each ordinary one-way ANOVA followed by Tukey’s multiple comparisons test with a single pooled variance, Pearson correlation analysis, unpaired two-tailed *t*-test, chi-square test, and two-tailed Mann–Whitney test was performed using GraphPad Prism v10.4.1 (https://www.graphpad.com).

## Acknowledgments

This work was supported by the University of British Columbia (UBC), the University of Northern British Columbia (UNBC) Office of Research, and the Natural Sciences and Engineering Research Council of Canada (NSERC) Discovery Grant 298521. This research was enabled in part by support provided by the British Columbia Digital Research Infrastructure Group and the Digital Research Alliance of Canada.

## Notes

### Competing Interest Statement

The authors have declared no competing interest.

